# Multimodal Single-Cell Omics Analysis of COVID-19 Sex Differences in Human Immune Systems

**DOI:** 10.1101/2020.12.01.407007

**Authors:** Yuan Hou, Yadi Zhou, Michaela U. Gack, Justin D. Lathia, Asha Kallianpur, Reena Mehra, Timothy Chan, Jae U. Jung, Lara Jehi, Charis Eng, Feixiong Cheng

**Author notes:** Correspondence to: Feixiong Cheng, PhD, Lerner Research Institute, Cleveland Clinic.

## Abstract

Sex differences in the risk of SARS-CoV-2 infection have been controversial and the underlying mechanisms of COVID-19 sexual dimorphism remain understudied. Here we inspected sex differences in SARS-CoV-2 positivity, hospitalization, admission to the intensive care unit (ICU), sera immune profiling, and two single-cell RNA-sequencing (snRNA-seq) profiles from nasal tissues and peripheral blood mononuclear cells (PBMCs) of COVID-19 patients with varying degrees of disease severity. Our propensity score-matching observations revealed that male individuals have a 29% increased likelihood of SARS-CoV-2 positivity, with a hazard ration (HR) 1.32 (95% confidence interval [CI] 1.18-1.48) for hospitalization and HR 1.51 (95% CI 1.24-1.84) for admission to ICU. Sera from male patients at hospital admission had decreased lymphocyte count and elevated inflammatory markers (C-reactive protein, procalcitonin, and neutrophils). We found that SARS-CoV-2 entry factors, including ACE2, TMPRSS2, FURIN and NRP1, have elevated expression in nasal squamous cells from males with moderate and severe COVID-19. Cell-cell network proximity analysis suggests possible epithelium-immune cell interactions and immune vulnerability underlying a higher mortality in males with COVID-19. Monocyte-elevated expression of Toll like receptor 7 (TLR7) and Bruton tyrosine kinase (BTK) is associated with severe outcomes in males with COVID-19. These findings provide basis for understanding immune responses underlying sex differences, and designing sex-specific targeted treatments and patient care for COVID-19.

## Introduction

Coronavirus Disease 2019 (COVID-19), which is caused by severe acute respiratory syndrome coronavirus 2 (SARS-COV-2), is a complex disorder with multisystem involvement across different organs^1–3^. SARS-CoV-2 has infected more than 9 million people and 253,769 people have died in the United States (US) since December, 2019 (Johns Hopkins data on November 20, 2020). Approximately 14% of COVID-19 positive patients show severe symptoms associated with advanced age^1^, sex^4^, genetics^5, 6^, disease comorbidities^1^, ^7^, and other risk factors. Yet, the mechanisms at the cellular and molecular level underlying these risk factors remain unclear, especially for sex differences impacting disease severity.

Sex differences in outcomes have been manifested in multiple infectious diseases, such as influenza^8^, hepatitis A and C^9^ viruses, and human immunodeficiency virus 1 [HIV1]^10, 11^. In addition, HIV1 and hepatitis C virus generally show a higher viral loads in men compared to women^12^. Moreover, women may mount higher immune responses to viral infections and vaccination^13^. An epidemiologic survey from 1965 through 1997 in the U.S. revealed that 80% of cases across 24 autoimmune diseases occurred in women^14^. Furthermore, in healthy populations, males have an elevated abundance of CD8= T cells, whereas women have higher proportions of CD4^=^ T cells and B cells in blood^15, 16^. These studies support the concept that sex differences in immune responses may play crucial roles in the incidence, progression, and outcomes of some human diseases, including COVID-19^4^.

During the COVID-19 pandemic, men have shown higher rates of critical cases, with a 6.6% increased mortality rate compare to women in the U.S. based on a report of 114,411 COVID-19-assosiatied deaths by the National Vital Statistics System from May 1 to August 31, 2020^17^. Similar observations were also made in United Kingdom, where males have a 1.78 hazard ratio of COVID-19-related deaths compared to females^1^. Moreover, male COVID-19 patients have a higher percentage of non-classical monocytes and elevated IL-8 and IL-18 levels in plasma^4^. Owing to heterogeneity of immune cells in the human body, the detailed genetic basis and molecular mechanism of sex differences that can explain the sex-specific risk of SARS-CoV-2 infection and disease severity remains unknown. Sex differences in immune responses in COVID-19 have a direct impact on the efficacy of vaccination and immune-related treatments. Hence, there is a pressing need to better understand the sex-specific heterogeneity of cell subpopulations of the human immune systems and its role in the severity of COVID-19.

In this study, we investigated sex differences in COVID-19 outcomes by combining observations from large-scale patient data from a COVID-19 registry and multimodal single-cell omics analysis of COVID-19 patient samples with varying degrees of disease severity. We identified that male patients have a higher susceptibility to severe COVID-19 using Propensity Score (PS)-adjusted observational analyses. By analysis of available laboratory testing data, we found that male patients had a lower level of circulating lymphocytes and elevated inflammatory markers (C-reactive protein, procalcitonin, and neutrophils) compared to female patients with COVID-19. We further performed multimodal single-cell omics data analysis of nasal tissues and peripheral blood mononuclear cells (PBMCs) isolated from COVID-19 patient blood to identify differential cell subpopulations contributing to sex differences of human immune responses. In summary, this study provides novel immunological mechanisms for the observed male bias in COVID-19 severity, which may offer individualized approaches for the prevention and treatment of male and female patients with COVID-19 in a sex-specific manner.

## Results

### Sex differences of COVID-19 outcomes influenced by age

In total, 27,659 individuals (8,361 COVID-19 positive) were tested between March 8 and July 27, 2020 within the Cleveland Clinic Health System in Ohio and Florida (**Table 1**). We observed that demographic factors (including age and race) were significantly different between females and males in the total cohort of the COVID-19 positive subgroup (**Table 1**). We found that female and male individuals have different percentage of comorbidities relevant to proven severity of COVID-19^1, 7^, including smoking (p < 0.001, two-tailed Fisher’s exact test), diabetes (p < 0.001), hypertension (p < 0.001), and coronary artery disease (p < 0.001). Interestingly, the fraction of COVID-19 positive females (n = 4,680, 56.0%) was higher than males (n = 3,681, 44.0%, p < 0.001, two-tailed Fisher’s exact test). We found that the sex difference in the occurrence of SARS-CoV-2 tested positive differed by age. For example, the prevalence of COVID-19 positivity greater in females than in males only in the age groups older than 80 years (80-90 years p = 0.006 and > 90 years p = 0.029, two-tailed Fisher’s exact test, **Supplementary Fig. 1a** and **Supplementary Table 1**). One possible explanation of overall high incidence of COVID-19 positive patients in female individuals (56.0%) compared to male individuals (44.0%) is that females have longer lifespan than males^18^.

**Table 1.** Cohort description with number of patients by sex in a COVID-19 registry.

|  | Total cohort |  |  | COVID-19 Positive |  |  |
| --- | --- | --- | --- | --- | --- | --- |
|  | Female | Male | p value | Female | Male | p value |
| **Patients (N)** | 16,354 | 11,305 |  | 4,680 | 3,681 |  |
| **Age (mean-SD)** | 48.9<br>(20.6) | 50.0<br>(21.0) | <0.001 | 49.6<br>(21.5) | 50.6<br>(19.6) | 0.029 |
| **White (%)** | 10,391<br>(63.5) | 7,080<br>(62.6) | 0.126 | 2,461<br>(52.6) | 2,019<br>(54.8) | 0.042 |
| **Black (%)** | 3,822<br>(23.4) | 2,489<br>(22.0) | 0.009 | 1,659<br>(35.4) | 1,157<br>(31.4) | <0.001 |
| **Race other (%)** | 828 (5.1) | 626 (5.5) | 0.087 | 245 (5.2) | 227 (6.2) | 0.074 |
| **Smoking (%)** | 1,766<br>(12.3) | 1,585<br>(16.6) | <0.001 | 279 (7.0) | 349<br>(11.7) | <0.001 |
| **COPD &**<br>**Emphysema (%)** | 1,217<br>(10.6) | 893<br>(11.7) | 0.016 | 286<br>(11.3) | 211<br>(10.8) | 0.682 |
| **Diabetes (%)** | 2,860<br>(23.8) | 2,429<br>(30.0) | <0.001 | 771<br>(28.5) | 769<br>(35.5) | <0.001 |
| **Hypertension (%)** | 6,032<br>(47.3) | 4,876<br>(55.8) | <0.001 | 1,787<br>(57.6) | 1,548<br>(62.4) | <0.001 |
| **Coronary artery**<br>**disease (%)** | 1,405<br>(12.1) | 1,722<br>(21.9) | <0.001 | 364<br>(14.3) | 448<br>(22.1) | <0.001 |
| **Hospitalization (%)** | - | - | - | 889<br>(19.0) | 957<br>(26.0) | <0.001 |
| **ICU admission (%)** | - | - | - | 245 (5.2) | 365 (9.9) | <0.001 |
| **ICU mechanical**<br>**ventilators (%)** | - | - | - | 99 (2.1) | 174 (4.7) | <0.001 |

We found that 26% of male patients (n = 957, p < 0.001, two-tailed Fisher’s exact test) compared to 19% (n=889) females were hospitalized for COVID-19; 9.9% (n = 365, p < 0.001, two-tailed Fisher’s exact test) of male patients compared to 5.2% (245) females were in intensive care units (ICU); and 4.7% (n=174, p < 0.001, two-tailed Fisher’s exact test) of male patients versus 2.1% (n=99) of females had to be mechanically ventilated in the ICU. Specifically, the male-predominant risks of hospitalization and ICU stay occurred in COVID-19 patients aged 50 to 90 years. The male predominance in COVID-19 disease severity is underscored further by the female predominance of COVID-19 infections over the age of 79 (**Supplementary Fig. 1a** and **Supplementary Table 1**).

### Sex is significantly associated with severe COVID-19 outcomes

We used an adjusted odds ratio (OR) to evaluate the associations between sex differences and COVID-19 outcomes after adjusting confounding factors using a propensity score matching approach. In total, we investigated four types of COVID-19 outcomes: (i) the SARS-CoV-2 positive rate by real-time reverse transcription polymerase chain reaction (RT-PCR), ii) hospitalization, iii) ICU admission, and (iv) whether the patients had to use mechanical ventilation in the ICU setting. To reduce risk of confounding factors, we adjusted for age, race, smoking, and four types of disease comorbidities (diabetes, hypertension, chronic obstructive pulmonary disease [COPD] & emphysema, and coronary artery disease) based on our sizeable efforts, using the PS matching method (see Methods). We found that male individuals were significantly associated with an increased likelihood of a positive laboratory test results by RT-PCR for SARS-CoV-2 (OR = 1.29, 95% confidence interval [CI] 1.18 – 1.41, **Fig. 1a**), COVID-19-related hospitalization (OR = 1.56, 95% CI 1.32 – 1.85), ICU admission (OR = 1.98, 95% CI 1.47 – 2.68), and requirement for mechanical ventilation (OR = 1.75, 95% CI 1.15 – 2.66) after adjusting for potential confounding factors. These observations together suggest that male individuals have an elevated incidence of SARS-CoV-2 infection and a higher likelihood of severe COVID-19 outcomes compared to females.

**Figure 1.**
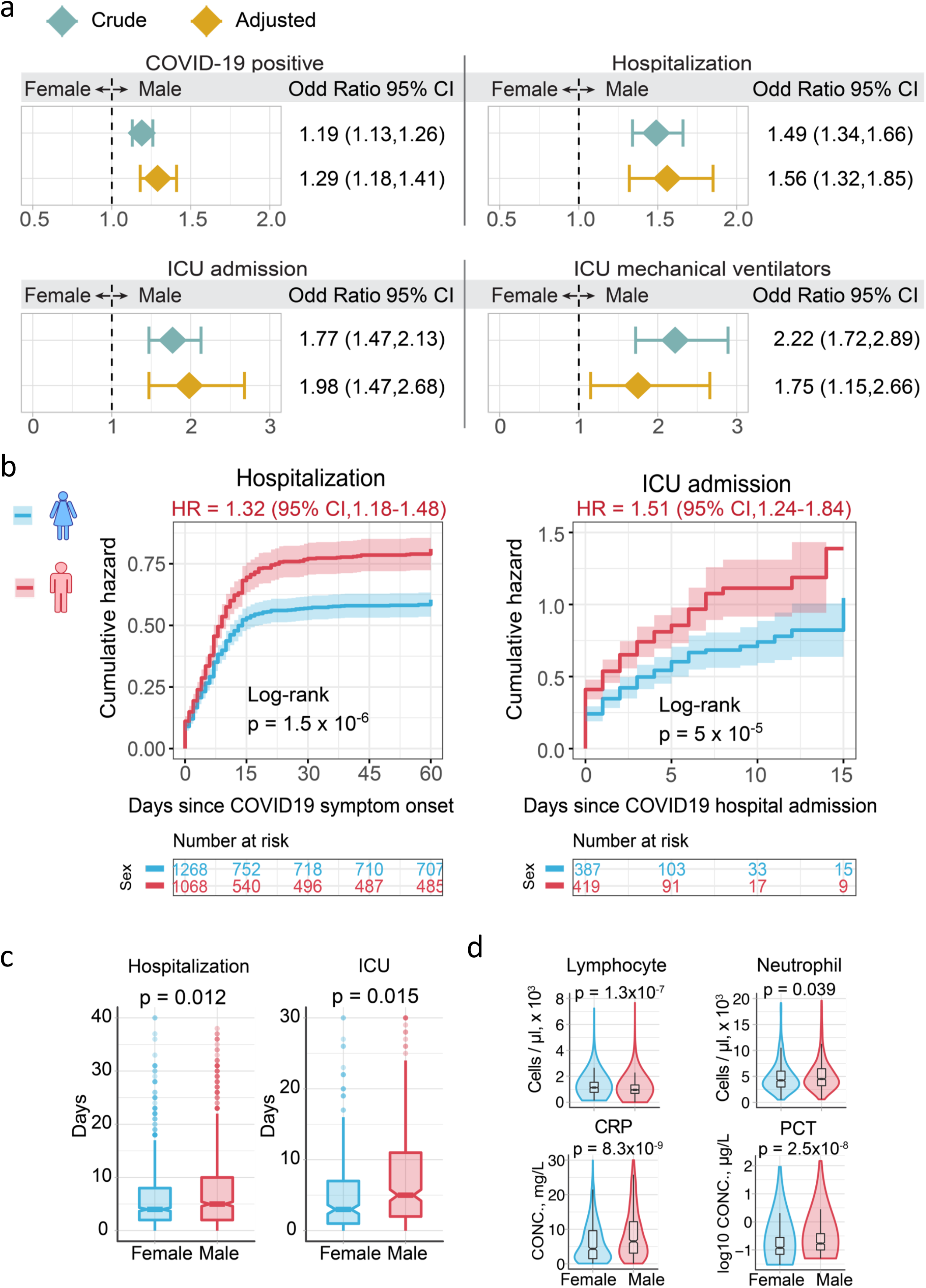
Male individuals are associated with severe COVID-19 outcomes. **a** Odds ratio (OR) analysis between sex and four COVID-19 outcomes: COVID-19 positive testing by RT-PCR, hospitalization, ICU admission, and usage of ICU mechanical ventilators. Crude cohort means that the OR value were computed based on original data. PS-adjusted OR ratio analysis: We used propensity score (PS) matching (1:1) population with similar covariate conditions (age, race, smoking, diabetes, hypertension, chronic obstructive pulmonary disease [COPD] & emphysema, and coronary artery disease; see Method). **b** Cumulative hazard of hospitalization and ICU admission. All results were computed in PS-matched groups. The log-rank test with the Benjamini & Hochberg (BH) adjustment are used to compare the statistical significance of cumulative hazard of hospitalization and ICU admission between males and females. The shadow represents 95% confidence interval. HR, hazard ratio, were computed using Cox proportional-hazards model. **c** Boxplots of the straying duration in hospital and ICU between male and female individuals. **d** Lab testing values for inflammatory markers between male and female individuals. P-value was computed by Kolmogorov– Smirnov test.

To better evaluate the hazard of sex differences on COVID-19 clinical outcomes, we performed Kaplan-Meier analysis to estimate the cumulative hazard between males and females for admission to hospital and ICU (**Fig. 1b and Supplementary Fig. 1b**). Male patients who tested positive for COVID-19 had a higher cumulative hazard for hospitalization than female patients using both PS-matching (hazard ratio (HR) = 1.32, 95% CI 1.18 – 1.48, p < 1.5 ×10^−6^, Log-rank, **Fig 1b**) and non-PS-matching methods (HR = 1.43, 95% CI 1.10 – 1.56, p < 1.9 ×10^−14^, Log-rank, **Supplementary Fig. 1b**). We found male patients had a longer duration of hospitalization than female patients (mean [±SD], 7.6 [±7.8] days versus [*vs.*] 6.2 [±6.7] days, p = 0.012, Kolmogorov–Smirnov [KS] test, **Fig 1c**). More specifically, males with COVID-19 also have a higher cumulative hazard for ICU admission compared to females (HR = 1.15, 95% CI 1.24-1.84, p < 2.0 ×10^−16^, PS-matching Log-rank test, **Fig 1b**). The average duration of ICU stays for male patients was 8.2 (SD = ±8.9) days, which is significantly longer than 6.2 (SD = ± 7.2) days for female patients (p = 0.015, **Fig 1c**). Altogether, our analysis suggests that male individuals are significantly associated with severe COVID-19 outcomes compared to female individuals.

### Sex-biased COVID-19 severity is associated with immune responses

Hyperinflammation has been reported as a major factor predisposing to a higher mortality in severe COVID-19 patients^19^ and there is a well-established sex difference in immune responses^13^. We next interrogated sex differences in inflammation-related clinical variables available in the COVID-19 registry. We found that the peripheral lymphocyte count was significantly lower in hospitalized male patients (p = 1.3×10^−7^, Kolmogorov–Smirnov [KS] test, **Fig 1d**) than in hospitalized females. In contrast, the circulating neutrophil levels in hospitalized male patients were higher than that of female patients (p = 0.039, **Fig 1d**). In addition, two other inflammatory parameters, C-reactive protein and procalcitonin, were significantly elevated in hospitalized males compare to females (p = 8.3×10^−9^ and 2.5×10^−8^, **Fig 1d**). Together, these observations reveal that the male-biased inflammatory responses are potentially associated with severe COVID-19. Yet, the underlying mechanisms of male-biased inflammatory responses for COVID-19 patients remain unknown. We therefore turned our attention to investigate this COVID-19 sexual-dimorphism in human immune systems using multimodal single-cell omics analysis.

### Sex-biased cell subpopulations in nasal tissues of critical COVID-19 cases

We investigated the single-cell RNA-sequencing (snRNA-seq) profiles of nasal tissues in the upper airway from COVID-19 positive patients *vs.* healthy donors (**Fig. 2a**). The samples comprised 11 critical COVID-19 patients, 8 moderate COVID-19 patients, and 5 healthy donors (**Supplementary Table 2**), as described in a previous study^20^. The snRNA-seq dataset contains 135,600 cells (**Fig. 2a**) across 22 annotated cell types within two main cell populations: epithelial cells (9 cell types) and immune cells (13 cell types, **Fig. 2b**).

**Figure 2.**
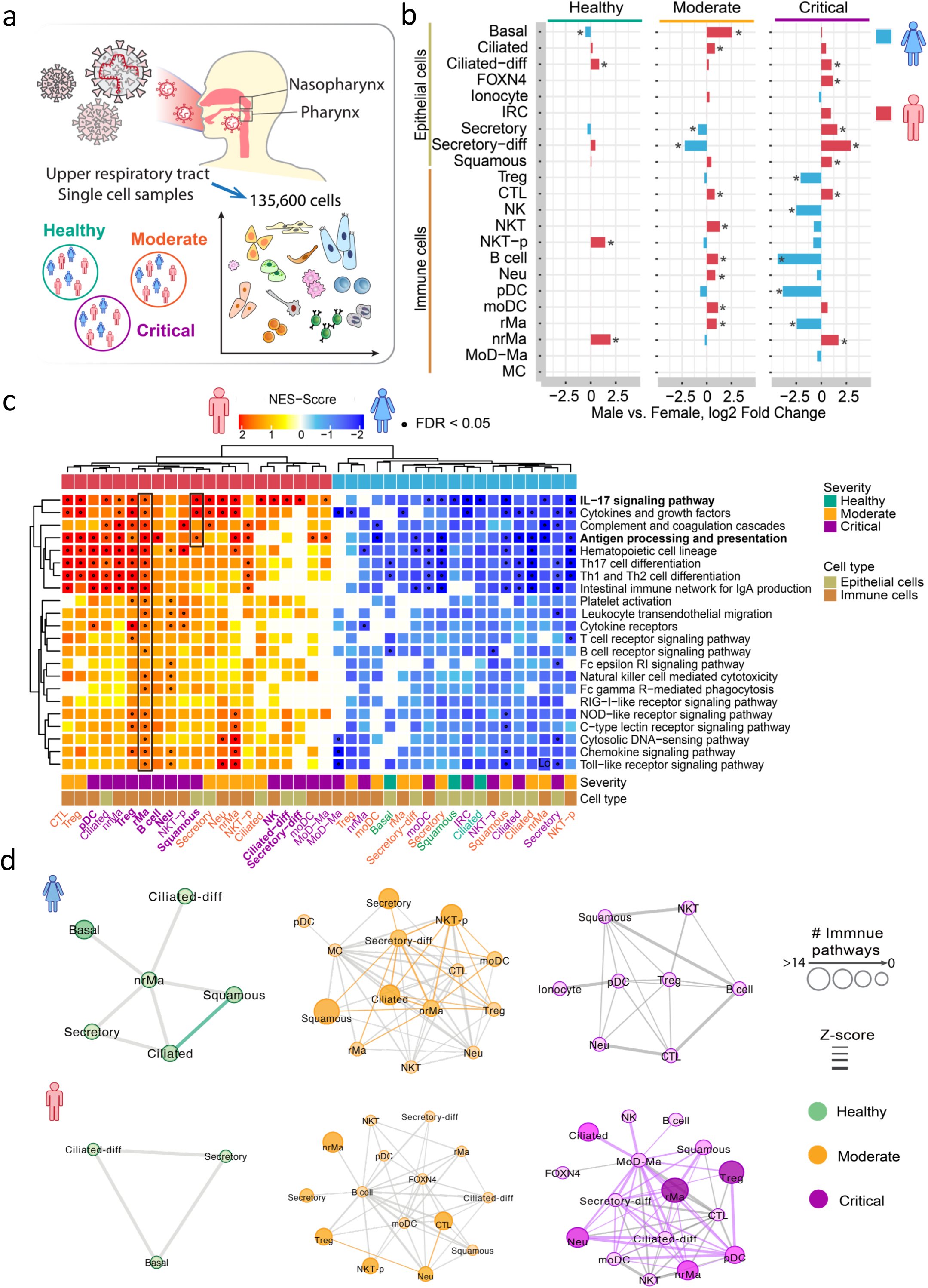
Sex-biased differential cell subpopulation and transcriptional analysis for the upper airway nasal tissues. **a** The graph show the sample information of single-cell RNA-sequencing analysis of nasopharynx and pharynx tissues by sex. **b** Bar-plots showing the log2 fold change of cell subpopulation abundances between male versus female across healthy donors, moderate and critical COVID-19 patients. Two-tailed Fisher’s exact test were conducted for each cell type by sex. *p < 0.05. **c** Gene-set enrichment analysis (GSEA) analysis of 22 immune pathways for male-biased gene set (up-regulated genes of male *vs*. female) and female-biased gene set (down-regulated genes of male *vs.* female, *see Methods*) across each cell types. The heatmap only showing the cell types with at least one significant immune pathway (FDR < 0.05). The 9 male-biased cell types in severe COVID-19 patients were highlighted by bold text. Black dots denote the FDR < 0.05 (The entire GSEA results are supply in **Supplementary Table 3**). **d** Sex-biased cell-cell interaction network analyses in healthy donors and COVID-19 patients. The weight of edge denotes the Z-score (Z < −2, p value < 0.001, *see Methods*) between two cells: more strong weight of edges means that sex-biased differentially expressed genes have close network proximity between two cell types in the human protein-protein interactome network. The size of nodes denotes the number of significantly enriched immune pathways by sex-based differentially expressed genes in a specific cell type. If the edge connected two immune activated cell types, it was shown the same color as nodes, otherwise it showed by gray.

From the analysis of relative proportions of each cell type, we observed that cell populations are significantly different between male and female patients with COVID-19 (**Fig. 2b**). Compared to the female patients with critical COVID-19, male patients with critical COVID-19 have a significantly elevated abundances across five epithelial cell types: the ciliated-diff (ciliated-differentiating, p < 2.0 × 10^−16^, Fisher test), FOXN4 (p = 6.1 × 10^−5^), secretory (p < 2.0 × 10^−16^), secretory-diff (secretory-differentiating, p < 2.0 × 10^−16^), and squamous cells (p < 2.0 × 10^−16^). For immune cells, male patients with critical COVID-19 have elevated abundances of CTL (Cytotoxic T cell, p < 2.0 × 10^−16^) and nrMa (non-resident macrophage, p < 2.0 × 10^−16^) compared to female patients (**Fig. 2b**). In contrast, the abundances of Treg (regulatory T cell, p < 2.0 × 10^−16^), NK (natural killer, p = 1.2 × 10^−7^), pDC (plasmacytoid dendritic cell, p < 2.0 × 10^−16^), and rMa (resident macrophage, p < 2.0 × 10^−16^) were decreased in critically ill male patients compared to critically ill females. These observations suggest that the differential epithelial and immune cell subpopulations between males and females may provide a possible explanation for the observed male-biased higher mortality in COVID-19 critical patients.

### Male-biased transcriptional networks in nasal tissues of critical COVID-19

We next sought to identify gene expression changes in sex-biased cell subpopulations from nasal tissues with varying degrees of COVID-19 severity. In each cell type, we defined the over-expressed genes between males *vs.* females (fold change [FC] > 1, FDR < 0.05, **Supplementary Fig 2a**) as the male-biased gene set; the female-biased gene set is the down-regulated genes in males *vs.* females (FC < 1, FDR < 0.05). We used the gene-set enrichment analysis (GSEA) to evaluate the 22 immune pathways at single-cell levels (**Supplementary Fig 2a**, see Method). We defined a cell type that has sex-biased differentially expressed genes enriched in at least one immune pathway (FDR > 0.05, **Supplementary Table 3**) as a sex-biased, immune perturbing cell type (**Fig 2c**). We found heterogeneous immune perturbing cell types for female patients across healthy donors, moderate and critical COVID-19 patients (**Fig. 2c**). Yet, we only found the elevated immune perturbing cell types for male patients with moderate and critical COVID-19 patients, but not for healthy donors. Compared to female patients, we identified 9 male-biased immune perturbing cell types for critical COVID-19, including squamous, ciliated-diff, secretory-diff, pDC, Treg, rMa, B cell, Neu and NK. Specifically, we found that 86% (19/22) of immune pathways were significantly enriched in sex-biased differentially expressed genes of rMa cells (FDR< 0.05) in male patients with critical COVID-19, including the IL-17 signaling pathway, cytokines and growth factors, antigen processing and presentation, Th17 cell differentiation, and Th1 and Th2 cell differentiation (**Fig. 2c**).

We further inspected network-based relationships of different immune cell types for sex-biased differentially expressed genes using network proximity measures, as described in our previous studies^21, 22^. We quantified network-based relationships using z-scores for sex-biased differentially expressed genes under the human protein-protein interactome network model for different immune cell types. We found spare cell-cell interaction networks for sex-biased differentially expressed genes in healthy donors between males and females (**Fig. 2d**). However, dense cell-cell interaction networks were observed for both male- and female-biased genes in moderate COVID-19 patients. Yet, we found much denser cell-cell interaction networks for male-biased differentially expressed genes in critical COVID-19 compared to female patients (**Fig. 2d**). In particular, we found that rMa and Treg were highly connected with several epithelial cell types, including squamous, secretory-diff, and ciliated-diff cell types, suggesting possible epithelium-immune cell interactions underlying sex differences of COVID-19. Taken together, both bioinformatics and network-based analysis revealed cell type-specific, male-biased transcriptional network activities for COVID-19 severity, such as epithelium-immune cell interactions.

### Activated squamous cells in males upon SARS-CoV-2 infection

Entry of SARS-CoV-2 into host cells depends on the expression level of the surface receptor ACE2 as well as S protein priming proteases^23^, including TMPRSS2^24^, FURIN^25^ and NPR1^26^. Yet, the expression levels of *ACE2* with either one of the S-priming proteases were unclear at single-cell levels between male and female individuals. We found that the epithelial cells had a higher expression of *ACE2* than immune cells in both males and females (**Supplementary Fig 3a**). We found that ACE2, as well as TMPRSS2 and FURIN, have an elevated expression levels in squamous cells from males than from female patients with both moderate and critical COVID-19 (**Fig 3a** and **Supplementary Fig 3b**). **Fig 3b** shows that *ACE2* is significantly co-expressed with *TMPRSS2* (p < 2.0×10^−16^), *FURIN* (p < 2.0×10^−16^) and *NPR1* (p = 2.3×10^−6^) in squamous cells from male patients with critical COVID-19 compared to female critical patients (**Fig 3b**).

**Fig. 3.**
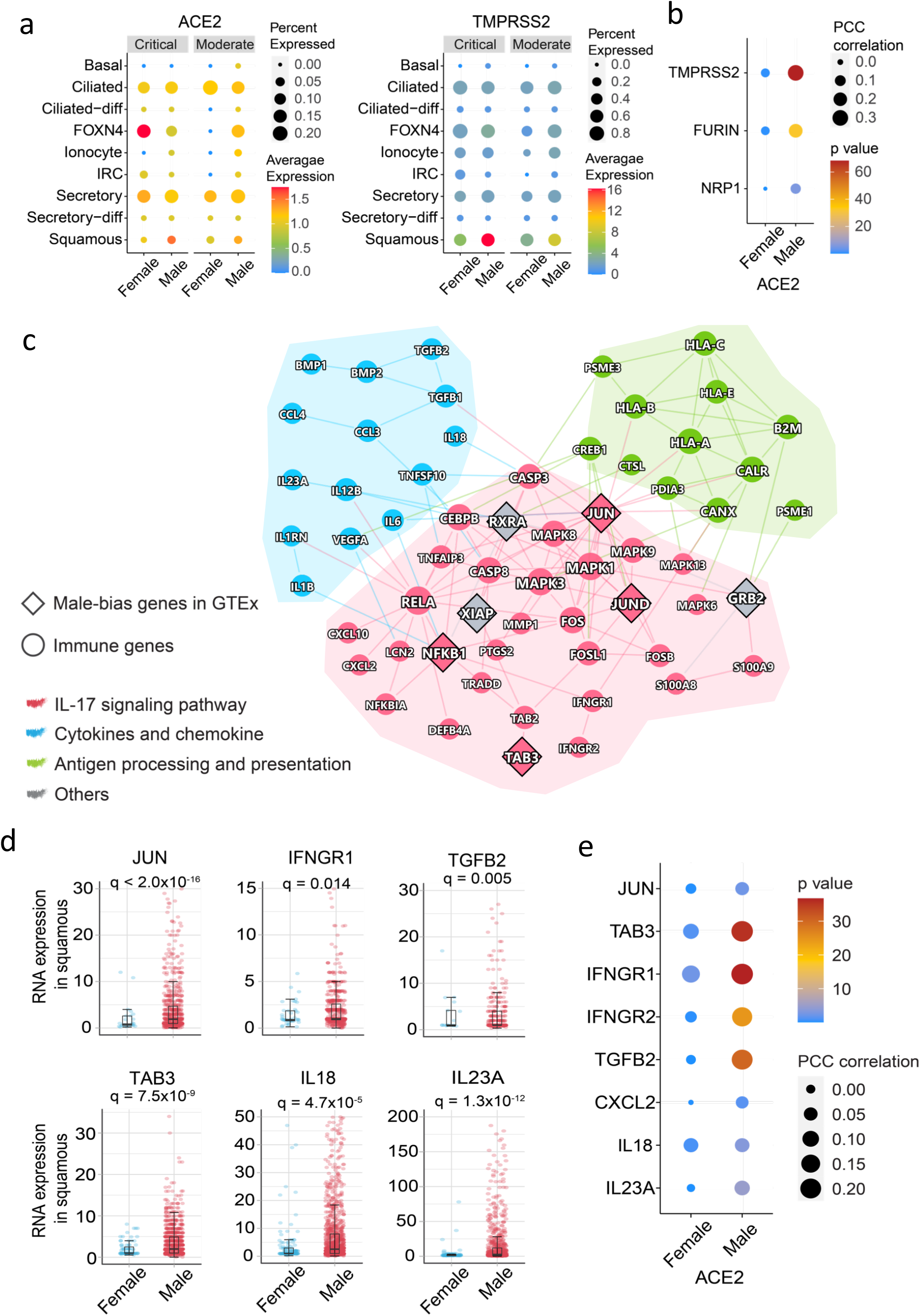
Male-biased transcriptional and network analysis of squamous cells of COVID-19 patients. **a** ACE2 and TMPRSS2 expression in epithelial cell types by sex. The size of dot denotes the percentage of ACE2 or TMPRSSE positive expressed cells. The gradient color bar represents the average expression of genes in each cell type. **b** Co-expression analysis of ACE2 with 3 S-protein priming proteases (TMPRSS2, FURIN, and NRP1). The size of dot denotes the Pearson Correlation Coefficient (PPC) values. The gradient color bar represents the p value (F-statistics) of PCC. **c** A highlighted protein-protein interaction subnetwork for male-biased differentially expressed immune genes in squamous cells from the patients with critical COVID-19. The colors for nodes and edges represents the different immune pathways. **d** The expression of selected male-biased genes of squamous cells from patients with critical COVID-19. Each dot means one cell, and the plot only show the gene positively expressed cells. For inside boxplots, the box represents the interquartile range (IQR). Adjusted p value (q) were computed by Benjamini-Hochberg method. **e** Co-expression dot plot of ACE2 with selected immune genes. The size of dot denotes the PPC values. The gradient color bar represents the p value of PCC.

We further performed GSEA analysis for male-biased gene set of squamous cells from patients with critical COVID19. We found that squamous cells from critically ill male patients were significantly enriched for 3 immune pathways (**Fig 2c**), including the IL-17 signaling pathway (FDR = 0.001), cytokines and growth factors (FDR = 0.001), as well as antigen processing and presentation (FDR = 0.047). We next turned to identify network modules (defined by the largest connect component [LCC] in the human interactome) for the male-biased immune gene set (up-regulated immune genes in males compared to females) of squamous cells. We found that male-biased immune genes significantly formed LCC (p = 0.04, permutation test, **Fig 3c**) in the human protein-protein interactome network. From the male-specific immune network of squamous cells, the sub-network of the IL-17 signaling pathway (red nodes) was highly connected to other immunological pathways. The proteins in the IL-17 signaling pathway were highly connected to the proteins in the antigen processing and presentation pathway through JUN, which was a male-biased transcription factor reported in Genotype-Tissue Expression (GTEx) database^27^. *TAB3* is an activator of JUN in the IL-17 signaling pathway, and the RNA expression levels of TAB3 (q = 7.5×10^−9^) is increased in squamous in male patients with critical COVID-19 (**Fig 3c-d**). We found that *TAB3* is an X chromosome-link (X-link) inactivated gene^27^, and 93% of *TAB3* expression are inactivated by XCI (X chromosome inactivation) in females^28^. These findings suggest that random X chromosome activation may explain some of the sex differences of COVID-19 disease severity in male and female individuals.

Notably, JUN and NFKB1 were enriched among male-biased genes in the GTEx database revealed by chromatin immunoprecipitation sequencing in promoter regions^27^. Compared to the female patients, *JUN* and *NFKB1* were highly expressed, with a broader distribution in squamous cells of male patients compared to females with critical COVID-19 (**Fig 3d and Supplementary Fig3c**). JUN and NFKB1 were found to induce the expression of multiple proinflammatory cytokines/chemokines and their receptors in male squamous, including *IFNGR1*, *IFNGR2*, *TGFB2*, *CXCL2* and *IL18* (**Fig 3c-d and Supplementary Fig 3c**). We also found that the increased proinflammatory cytokines (*IFNGR1*, *IFNGR2* and *TGFB2*) were significantly co-expressed with *ACE2* in male squamous cells from critically ill COVID-19 patients (**Fig 3e**). Altogether, these observations suggest that squamous cells play crucial roles in male-biased mortality for critical COVID-19 patients. Further independent cohort validation and functional observations are highly warranted.

### Male-biased immune cell subpopulations in PBMCs from severe COVID-19 cases

To understand the sex-specific host immune responses at the single-cell level, we utilized PBMC single-cell RNA-sequencing datasets from COVID-19 patients (n = 9) and healthy donors (n = 4, **Fig. 4a** and **Supplementary Table 2**). In total, we re-analyzed 49,054 cells and clustered them to 13 annotated cell types based on well-defined maker genes (**Supplementary Fig 4**) and 3 un-annotated cell types (see Methods) from a recent study (see Methods).

**Fig. 4.**
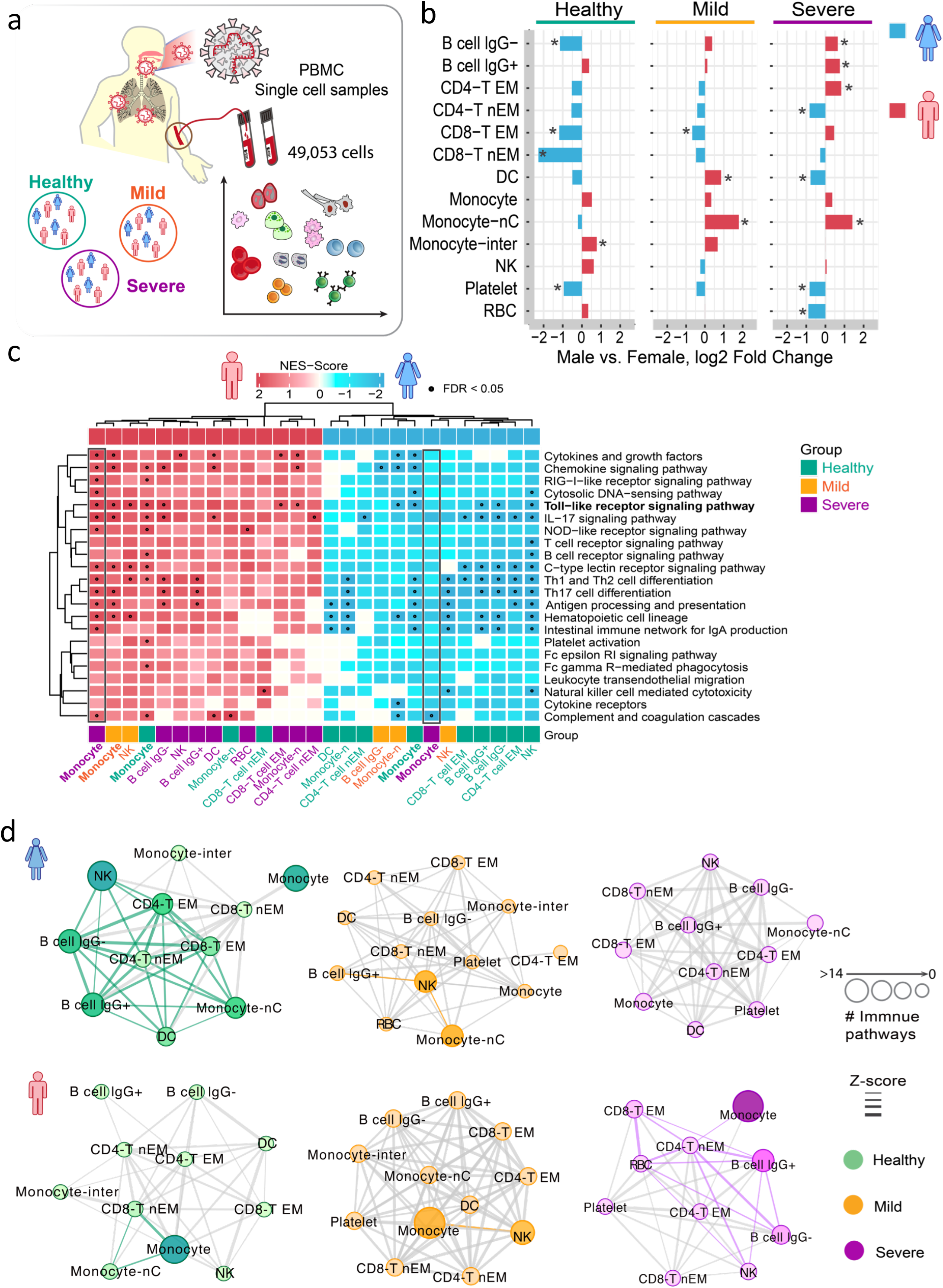
Sex-biased differential cell subpopulation and transcriptional analysis for peripheral blood mononuclear cells (PBMCs). **a** A diagram show the information of single cell RNA-sequencing analysis of PBMC by sex. **b** Bar-plots showing the log2 fold change of male versus female in cell type abundances of PBMCs isolated from bloods of healthy donors, and patients with mild or severe COVID-19. Two-tailed Fisher’s exact test were conducted for each cell type by sex. *p < 0.05. **c** Gene-set enrichment analysis (GSEA) analysis of 22 immune pathways for male-biased gene set (up-regulated genes of male *vs.* female) and female-biased gene set (down-regulated genes of male *vs*. female, see Methods) across each cell types. The heatmap only showing the cell types with at least one significant immune pathway. Black dots denote the FDR < 0.05 (The entire GSEA results are supply in **Supplementary Table 4)**. **d** Sex-biased cell-cell interaction network analyses in healthy donors and COVID-19 patients. The weight of edge denotes the Z-score between two cells (z < −2): more strong weight of edges means that sex-biased differentially expressed genes have close network proximity between two cell types in the human protein-protein interactome network. The size of nodes denotes the number of significantly enriched immune pathways by sex-based differentially expressed genes in a specific cell type. If the edge connected two immune activated cell types, it was shown the same color as nodes, otherwise it showed by gray.

We identified multiple, sex-specific differential immune cell types from PBMCs across healthy controls, mild and severe COVID-19 patients (**Fig 2b and 4b**). Compared to female COVID-19 patients, the male patients with severe and mild COVID-19 had significantly elevated abundances of monocytes-nC (non-classic monocytes, p < 2.0 × 10^−16^ [mild] and p = 3.7 × 10^−5^ [severe], **Fig 4b**). In parallel, we found that male patients in the severe COVID-19 group have higher abundances of B cells (lgG-[lgG non-expressed B cell], p = 4.7 × 10^−11^ and lgG= [lgG expressed B cell], p = 1.7 × 10^−9^) and CD4-T EM (effector memory like CD4 T cells, p = 1.3 × 10^−9^) but lower abundances of CD4-T nEM cells (non-effector memory like CD4 T cells, p = 1.1 × 10^−7^) and DC cells (p = 0.016).

We further used the GSEA method to evaluate the immune pathway activities based on male/female-biased gene sets identified from snRNA-seq data of PBMCs (**Supplementary Fig 2a** and **Supplementary Table 4**). We found that more cell types were significantly enriched by immune pathways in PBMCs from male patients with severe COVID-19 compared to females (**Fig. 4c**). Yet, fewer cell types were enriched by immune pathways in PBMCs of male healthy donors compared to healthy females (DC, monocyte, monocyte-nC, CD4-T cell nEM/EM, CD8-T cell EM, B cell lgG-/= and NK). These results suggest that male patients may have more significant immune responses than females after SARS-CoV-2 infection. It should be pointed out that 14 immune pathways were significantly enriched in monocytes (FDR< 0.05) from male patients with severe COVID-19 (**Fig. 4c**), such as Toll-like receptor signaling pathways, RIG-I-like receptor signaling pathway, cytokines and growth factors, and the IL-17 signaling pathway. Yet, in female severe COVID-19 patients, only the complement and coagulation pathways were significantly enriched in the monocytes (FDR = 0.003).

We next inspected the cell-cell network relationships of PBMCs from both male and female patients across healthy controls, mild and severe COVID-19 patients. Using network proximity measure, we found a similar cell-cell interaction network for female patients with mild and severe COVID-19 (**Fig 4d** and **Supplementary Fig 5a**). Yet, male patients with severe COVID-19 significantly have the elevated immune cell-cell interactions compared to males with mild COVID-19, revealing possible immune vulnerability associated with male-biased high mortality in severe COVID-19. For example, monocyte-activated immune responses were predominantly observed in cell-cell interaction networks of male-derived PBMCs with severe COVID-19 compared to female patients (**Fig 4d**). In summary, both bioinformatics and network analyses suggest that immune vulnerability may be associated with male-biased morbidity and mortality in severe COVID-19, such as elevated monocyte-related immune responses.

### Elevated monocyte immune responses in male patients with severe COVID-19

We further performed human protein-protein interactome network analysis to investigate the immune characteristics of monocytes in male patients with severe COVID-19. We found that the male-biased gene set (up-regulated genes in male monocytes *vs.* female monocytes) formed the significant network module (**Fig 5a**, p < 0.001, Permutation test) in the human interactome. This male-biased, monocyte-specific protein-protein interaction network was significantly enriched by several key immune pathways, including Toll-like receptor pathway (gold), IL-17 signaling pathway (red), cytokines and growth factors (blue), and antigen processing and presentation (green) (**Fig 5a**). Several hub genes (including *JUN, NFKB1, CCR1* and *SATA1*) are highly connected among different immune pathways. Specifically, the expression level of JUN and NFKB1 in monocytes is significantly increased in male patients with severe COVID-19 patients compared to females (q < 2.0 × 10^−16^).

**Fig. 5.**
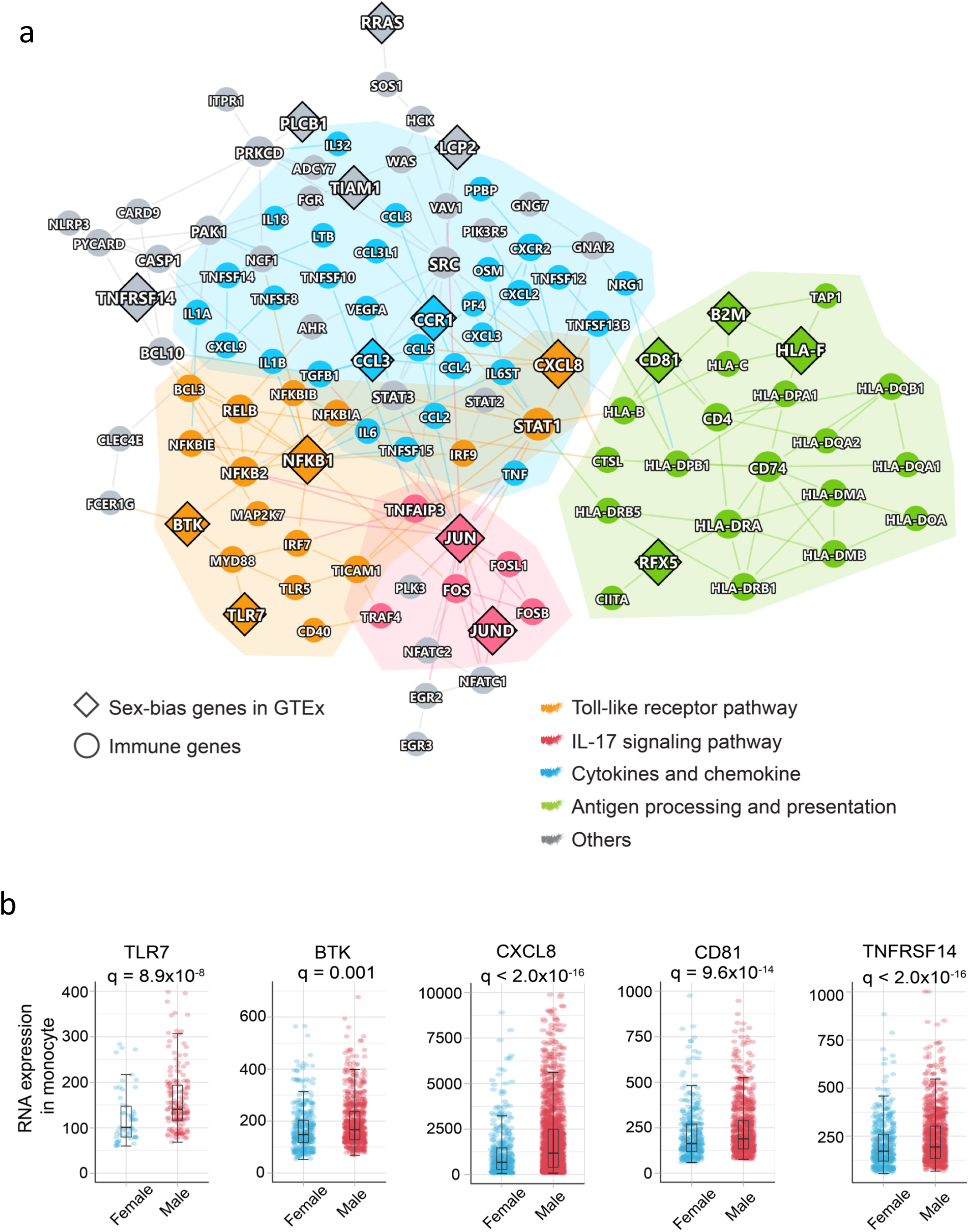
Elevated monocyte immune responses in male patients with severe COVID-19. **a** A highlighted protein-protein interaction subnetwork for male-biased differentially expressed immune genes in monocytes from the patients with severe COVID-19. The colors for nodes and edges are represents the different immune pathways. **b** The expression of selected male-biased immune genes in monocytes from the patients with severe COVID-19. Each dot demotes one cell, and the plot only show the gene positively expressed cells. For inside boxplots, the box represents the interquartile range (IQR). Adjusted p value (q) were computed by Benjamini-Hochberg method.

We next selected genes using subject matter expertise based on a combination of factors: a) sex-biased expression in GETx, b) available eQTL data, c) well-annotated immune genes from KEGG database^29^, and d) available evidences of X-link genes. Applying these criteria resulted in two top-predicted sex-based genes, including Toll-like receptor 7 (*TLR7*) and Bruton tyrosine kinase *(BTK),* which may explain the disease severity of COVID-19. Both *TLR7* and *BTK* are X-link inactive genes. Specifically, 84% of *TLR7* and 98% *BTK* expression are inactivated by XCI in females^28^. A recently study found that *BTK* was associated with one SNP rs2071223 in male lymphocytes (p = 0.005)^27^. In a case series of 4 young men with severe COVID-19, rare putative loss-of-function variant (c.2129_2132del) of TLR7 were reported to be associated with impaired type I and II IFN responses^30^. We found that *TLR7* (q = 8.9×10^−8^) and *BTK* expression (q = 0.001) in monocytes are significantly increased in male patients with severe COVID-19 compared to females (**Fig. 5b**). Furthermore, the downstream factors of TLR7 and BTK in Toll-like receptor pathway, such as MYD88, IRF7, NFKB1, JUN, and cytokines TNF, IL1B, IL18 were also found to be significantly increased in male patients with severe COVID-19 (**Fig. 5b and Supplementary Fig 5b**). Altogether, monocyte-specific expression of *TLR7* and *BTK* may provide potential explanations for the male-biased disease severity in COVID-19.

## Discussion

In this study, we comprehensively investigated the sex differences in disease severity and mortality between male and female individuals using an integrative approach that integrates large-scale COVID-19 patient registry, multimodal analysis of snRNA-seq profiles, and human protein-protein interactome network analysis. We identified that male patients with COVID-19 had a higher rate of hospitalization and ICU admission, and a longer stay time in hospital or ICU, compared to female individuals (**Fig. 1**). Our finding is consistent with an observational study using 17 million COVID-19 tested populations: males individuals had 1.78 hazard for COVID-19 related death compared to female individuals^1^.

Via analysis of laboratory testing data in the COVID-19 patient registry database, we found that serum of male patients has elevated inflammatory markers (C-reactive protein, procalcitonin, and neutrophil) compared to female patients with COVID-19, suggesting sex-specific immune responses underlying sex differences between male and female individuals with COVID-19. We further performed multimodal analysis of single-cell RNA-sequencing data from both nasal tissues and PBMCs with varying degrees of COVID-19 pathology. We identified several sex-biased, differential immune cell types and gene transcriptional networks that may provide explanations for the higher mortality in male patients with severe COVID-19. For example, network proximity analysis of sex-biased, differentially expressed genes in the human protein-protein interactome show potential epithelium-immune cell interactions from nasal tissues and immune vulnerability from PBMCs underlying male-biased high mortality for COVID-19.

ACE2, a key SARS-CoV-2 receptor in host cells^23, 31^, together with the S-protein priming proteases TMPRSS2, FURIN and NPR1, facilitate viral entry into the human upper respiratory tract^24–26^. High expression levels of ACE2, TMPRSS2 and FURIN is predicted to results in an enhanced efficiency of SARS-CoV-2 infection and greater severity of COVID-19^20^. Aligned with the observation of COVID-19 severity, we found that several SARS-CoV-2 entry factors, including *ACE2, TMPRSS2, FURIN* and *NRP1,* have elevated expression in nasal squamous cells of male individuals with both moderate and severe COVID-19, but not in females (**Fig 3** and **Supplementary Fig 3b**). This finding may provide a possible explanation for the 29% increased likelihood of SARS-CoV-2 infection in men *vs.* women in our COVID-19 registry (**Fig. 1a**). In addition, we found that *ACE2* was significantly co-expressed with multiple innate immune genes (including *JUN, IFNGR1, TGFB2, CXCL2, IL18,* and *IL23A*) in nasal/respiratory squamous cells from male patients with severe COVID-19. All these findings imply that sex-biased, differential epithelium cell subpopulations (**Figs 2b** and **4b**) and transcriptional network changes (**Figs 2d** and **4d**) may contribute to higher incidence in male patients with severe COVID-19 compared to female individuals.

A recent study showed that the myeloid cell dysregulation was a marker for severe COVID-19 diaease^32, 33^. In our current study, we found that differential transcriptomes in the rMa of nasal tissue and PBMCs were significantly enriched in multiple immune pathways (**Fig. 2c** and **Fig. 4c**) in male patients with severe COVID-19. These findings provide a potential explanation as so why that male COVID-19 patients often have a stronger likelihood of increased morbidity (1.32 HR in hospital and 1.51 HR in ICU, **Fig1 b**) and a higher mortality than female patients.

TLR7 and BTK are male-biased, differentially expressed genes in peripheral monocytes of male patients with severe COVID-19. TLR7 escapes the XCI in female B cell, resulting in the higher expression levels in women than in men^34^. Young men with severe COVID-19 were found to carry the 4-nucleotide deletion in *TLR7* (c.2129_2132del; pGln710Argfs*18), while the affected family members carried only one missense variant on *TLR7* (c.2383G>T; pVal795Phe)^30^. This evidence supports the potential role of TLR7 in the male-associated higher mortality seen in severe COVID-19 patients (**Fig. 5b**). BTK, a tyrosine kinase, was identified as a top male-biased, differentially expressed gene in monocytes of severe COVID-19 patients. BTK is an X-linked gene and 98% of its expression is inactivated by XCI^28^ in females. By analyzing gene expression profiles of 838 subjects from the GETx database, we found that an eQTL SNP (rs2071223) on BTK in male-derived lymphocytes (p = 0.005)^27^ but not female, further supporting the male-specific role of BTK in COVID-19. Several BTK inhibitors (*e.g.,* acalabrutinib and ibruitinib, which blocks TLR7-dependent NF-κB activation in monocytes), have been shown to be potentially promising in treatment of patients with severe COVID-19. Altogether, these observations emphasize that sex is a key biological variable which should be considered in predicting the efficacy of pharmacologic treatments (such as BTK inhibitors) in people diagnosed with COVID-19.

We acknowledge several potential limitations of our study. Samples sizes of snRNA-seq datasets analyzed in this study are relatively small; and the smaller number of female patients compared to males may influence the findings of differential cell subpopulation analysis. Thus, the sex-biased cell types and transcriptional networks we identified should be validated further in prospective large-scale cohorts with varying degrees of COVID-19 pathology, including asymptomatic patients. In addition, we observed that the male patients aged between 30 to 80 years have a greater risk of infection by SARS-CoV-2 (**Supplementary Fig 1a**), but in the group of females aged 80 years or older females had a higher prevalence of confirmed SARS-COV-2 infections. Exploring the sex differences and underlying immune mechanisms in younger COVID-19 patients, including the pediatric population, may provide more actionable biomarkers and immune targets for disease prevention and vaccine development^35, 36^. Finally, genetic basis of sex differences should be investigated in the future using the genetic datasets from the growing COVID-19 population, such as the genome-wide association studies from COVID-19 Host Genetics Initiative^37^ (https://www.covid19hg.org/).

Taken together, our analysis provides a comprehensive understanding of the clinical characteristics and immune mechanisms underlying sex differences in COVID-19. We found that male individuals with COVID-19 have significantly elevated rates of hospitalization and ICU admission, and longer stay times in hospital or ICU. Laboratory testing variable observation, bioinformatics and network-based analyses of single-cell data from nasal tissues and PBMCs provide possible immunological explanations for the identified male-biased severity and higher mortality in COVID-19. If broadly applied, these findings will offer a path toward sex-specific molecularly targeted therapeutic development for COVID-19, which will be essential against the COVID-19 pandemic and future pandemics from other emerging pathogens.

## Methods and Materials

### COVID-19 registry

We used the institutional review board–approved COVID-19 registry data, including 27,659 individuals (8,274 positive) tested during March to July, 2020 from the Cleveland Clinic Health System in Florida and Ohio. All tested samples were pooled nasopharyngeal and oropharyngeal swab specimens. Then the infection with SARS-CoV-2 was confirmed by RT-PCR in the Cleveland Clinic Robert J. Tomsich Pathology and Laboratory Medicine Institute. All SARS-CoV-2 testing was authorized by the Food and Drug Administration under an Emergency Use Authorization and accord with the guidelines established by the Centers for Disease Control and Prevention.

The data in COVID-19 registry include COVID-19 test results, baseline demographic information, medications, and all recorded disease conditions and others. We conducted a series of retrospective studies to test the sex difference with four COVID-19 outcomes, COVID-19 test positive. Data were extracted from electronic health records (EPIC Systems) and were manually checked by a study team trained on uniform sources for the study variables. We collected and managed all patient data using REDCap electronic data capture tools. The statistical analysis for smoking, diabetes, hypertension, chronic obstructive pulmonary disease [COPD] & emphysema and coronary artery disease were calculated after missing value deletion.

### Propensity score (PS) matching analysis

We select case-control propensity score (PS) method to matching our four COVID-19 outcomes: i) COVID-19 positive outcome: COVID-19 positive patients (n = 8361) were matched to negative patients (n = 19,298) in total COVID-19 testing patients; ii) Hospitalization outcome: hospitalized patients (n = 1,846) were matched to non-hospitalized patients (n = 6,515) in COVID-19 positive patients; iii) ICU admission outcome: the patients of ICU admission (n = 610) were matched to non-ICU admission patients (n = 1,236) but they were in hospital due to COVID-19; iv) ICU mechanical ventilator: the COVID-19 patients used mechanical ventilator (n = 273) were matched to non-mechanical ventilator user (n = 337) in ICU. To reduce the bias from confounding factors, all PS-matched patients were adjusted for age, race, smoking, presence of diabetes, hypertension, chronic obstructive pulmonary disease [COPD] & emphysema and coronary artery disease. The PS-matched groups were used to the next clinical analysis. The PS-matching analyses were conducted by matchit package in the R v3.6.3 platform.

### Clinical outcome analysis

The odds ratio (OR) was used to measure the association between the COVDI-19 outcomes and sex based on logistic regression model. An OR > 1 means that male sex is associated with a higher likelihood of the outcome. The Kaplan-Meier method was used to estimate cumulative hazard of hospitalization and ICU-admission of COVID-19 positive patients by sex. And Cox proportional regression model was used to quantified the hazard of sex for COVID-19 outcomes. For hospitalization outcome, the time was calculated from the start date of COVID-19 symptom onset to hospital admission date. For ICU admission outcome, the time was calculated from the date of patients admitted to hospital to the date of ICU admission. Log-rank test was used for comparison among different sex with BH (Benjamini & Hochberg) adjustment^38^. All the cumulative hazard analyses were performed using the Survival and Survminer packages in R 3.6.0 (https://www.r-project.org).

### Single-cell RNA-seq data analysis

In this study we used two single-cell datasets of COVID-19 patients versus healthy control^20^ (**Supplementary Table 2)**. i) Dataset-1 (European Genome-phenome Archive repository: EGAS00001004481), it was collected nasopharyngeal and pharyngeal tissues from COVID-19 positive patients (11 severe patients with 3:8 female *vs.* male ratio, and 8 mild patients with 1:7 female *vs.* male ratio) and healthy control (5 healthy donors with 3:2 female *vs.* male ratio). The dataset contains 135,600 cells with cell type annotated. Therefore, all analysis in dataset-1 were based on their cell type annotation. ii) Dataset-2 (GSE149689)^39^ was downloaded from the NCBI GEO database. This data set included three groups, patients infected influenza A, COVID-19 and healthy controls. But we only focus on COVID-19 and healthy control population in this study. For the COVID-19 group, the peripheral blood mononuclear cells (PBMC) samples were collected from 4 severe patients (1:1 female *vs.* male ratio) and 5 mild patients (3:2 female *vs*. male ratio). In addition, 4 donors in healthy control group with 3:1 ratio in female *vs.* male. In total, 49,053 cells were used to next analysis. Qualifying cells based on the criteria from the original paper were used for the single cell analysis. We used the cell type gene markers from a previous study^39^ (CD3E, CD4, CCR7, CD8A, NCAM1, CD14, FCGR3A, NR4A1, CD19, FCER1A, PPBP and HBB, **Supplementary Fig. 4b**). All single-cell data analyses and visualizations were performed with the R package Seurat v3.1.4 ^40^. “NormalizeData” was used to normalize the data. “FindIntegrationAnchors” and “IntegrateData” functions were used to integrate cells from different samples. tSNE was used as the dimension reduction method for visualization. ‘FindAllMarkers’ function with the MAST test as the finding maker method for each cell type.

### Sex specific differences in gene expression by cell types

The sex specific expressed gene sets were defined as the differentially expressed genes in each cell type of of male *vs.* female. edgeR^41^ v 3.12 was used to computing the differential expression genes in different cell types based on R platform 4.0. Male specific gene sets of each cell type were defined as significantly up-regulated genes of males *vs.* females (log2 fold change > 0 and FDR < 0.05, **Supplementary Fig 2a**), and female specific gene sets in each cell type were defined as the significantly down-regulated genes in males *vs.* females (log2 fold change < 0 and FDR < 0.05).

### Immune gene set enrichment analysis

To evaluated the immune pathway activity in female and male, GSEA was conducted as described in previous work^42^. The immune gene profiles were retrieved from KEGG database^29^. We selected 22 immune related pathways and 1241 genes from KEGG belonging to the immune system subtype. The normalized enrichment score (NES, Equation 1) was calculated for 22 immune pathways in male and female specific gene sets (**Supplementary Fig 2a**). The equation is shown as follows:

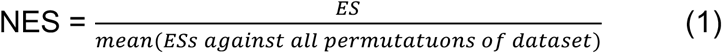

where ES^42^ denotes enrichment score. Through normalizing the enrichment score, GSEA avoid the differences in gene set size and in correlations between gene sets and the expression dataset. For male gene sets of this study, we only selected NES > 0 and FDR < 0.05 pathways as male activated immune pathways. For female gene sets, we only selected NES < 0 and FDR < 0.05 pathways as the female activated immune pathways. Permutation test (1000 times) was performed to evaluate the significance. All analyses were performed with the prerank function in GSEApy package (https://gseapy.readthedocs.io/en/master/index.html) on Python 3.7 platform.

### Functional enrichment analysis

We performed KEGG enrichment analyses to reveal the biological relevance and functional pathways. All functional enrichment analyses were performed using Enrichr^43^. And the FDR < 0.05 as significantly enriched pathways.

### Building the human protein-protein interactome

To build a comprehensive human interactome, we assembled in total 18 bioinformatics and systems biology databases to collect PPIs with five types of experimental evidences: (1) literature-curated PPIs identified by affinity purification followed by mass spectrometry (AP-MS), Y2H, literature-derived low-throughput experiments, or protein three-dimensional structures from BioGRID^44^,IntAct^45^, Instruct^46^, MINT^47^, PINA v2.0^48^ and InnateDB^49^; (2) binary PPIs tested by high-throughput yeast-two-hybrid (Y2H) systems from two public available high-quality Y2H datasets^50, 51^ and one in-house dataset^52^; (3) kinase-substrate interactions by literature-derived low-throughput or high-throughput experiments from Kinome NetworkX^53^, Human Protein Resource Database (HPRD)^54^, PhosphositePlus^55^, PhosphoNetworks^56^, Phospho.ELM^57^ and DbPTM 3.0^58^; (4) signaling network by literature-derived low-throughput experiments from SignaLink 2.0^59^; and (5) protein complexes data identified by a robust affinity purification-mass spectrometry methodology collected from BioPlex v2.0^60^. The final human protein-protein interactome used in this study included 351,444 unique PPIs (edges or links) connecting 17,706 proteins (nodes). The detailed description for building human protein-protein interactome are provided in our recent studies^21, 22, 61^.

### Cell-cell proximity measure

We used the “shortest” network proximity metric to evaluate the cell-cell interactions in male and female. First, we generated the sex specific gene set, which defined as significant differential expression genes by male *vs.* female in each cell type (FDR < 0.05, **Supplementary Fig 2a**). Since the sizes of the gene sets vary largely, we selected the top 200 differentially expressed genes based on the fold change for the gene sets that have more than 200 genes. Next, for two gene sets *A* and *B*, their “shortest” network proximity *d*_*AB*_ was calculated as:

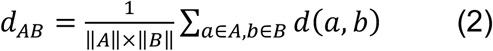

where *d*(*a*, *b*) is the shortest distance of *a* and *b* in the human interactome. To evaluate the significance of the proximity, we performed a permutation test repeated 1,000 times using semi-randomly selected genes/proteins that have similar degree distributions to the two gene sets being evaluated. We then calculated the Z score as:

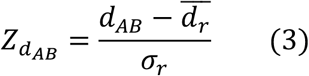

where 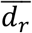 and *σ*_0_ were the mean and standard deviation of the permutation test.

### Identification of cell type-specific, sex-biased immune gene networks

We picked the overlap genes between sex specific differential genes set and 1241 immune genes (22 immune pathway from KEGG) as sex-biased immune gene set for each cell type. In addition, we identified some immune genes as highly confidential sex-bias genes based on the following criteria: 1) the genes were X-chromosome linked genes from GTEx and public literature. 2) Sex-biased transcription factors and other genes in specific tissues or cell types from GTEx database. 3) the genes were significantly associated with sex-biased eQTL in specific tissues or cell types (the solid tissues which majority cell type are epithelial cell, blood and lymphocytes). Thereafter, we picked largest connected component from sex-biased immune gene set based on PPIs as final sex-biased immune gene module in specific cell type. This step was performed with the NetworkX package (https://networkx.github.io/) on Python 3.7 platform.

### Statistical analysis and network visualization

Statistical tests for assessing categorical data through *χ*^2^ was performed by SciPy 1.2.1 (https://www.scipy.org/). The one-way ANOVA was used to compare the difference of continuous clinical variable by sex. All statistical analyses with the significance level set at p < 0.05 were used. Networks were visualized using Cytoscape.

## Author contributions

F.C. conceived the study. Y.H. and Y.Z. performed all experiments and data analysis. M.U.G., A.K., J.D.L., R.M., T.C., J.U.J., L.J., and C.E., discussed and interpreted results. Y.H. and F.C. wrote and critically revised the manuscript with contributions from other co-authors.

## Data availability statement

All codes and data used in this study are free available: https://github.com/ChengF-Lab/COVID-19Sex. Other data are available in Supplementary file and other codes used in this study are available upon reasonable correspondence to the corresponding authors.

## Supporting Information

**Supplementary Fig.1 Clinical outcome and characteristics between male and female individuals with COVID-19. a** Statistics analysis of four COVID-19 outcomes across different age groups. * denote p < 0.05 using two-tailed Fisher’s extract test. **b** Cumulative hazard of hospitalization and ICU admission are shown. The log-rank test with the Benjamini & Hochberg (BH) adjustment was used for comparing the statistical significance of cumulative hazard of hospitalization and ICU admission between men and women. The shadow represents 95% confidence interval. HR, hazard ratio.

**Supplementary Fig.2** Cell types analysis of nasal samples by sex. **a** Workflow of GSEA analysis. **b** Heatmap showed the z-score in male and female patients with critical COVID-19.

**Supplementary Fig.3** The expression of ACE2 and immune genes by sex. **a** ACE2 expression by sex in 22 cell types across critical and moderate COVID-19. conditions. The size of dot denotes the percentage of ACE2 or TMPRSSE positive expressed cells. The gradient color bar represents the average expression of genes in each cell type. **b** the dot plot showed the expression level and distribution of FURIN and NPR1 in epithelial cells by sex. **c** The expression of male-biased immune genes of squamous in the patients with critical COVID-19. Each dot means one cell, and the plot only show the genes positive expressed cells. For inside boxplots, the box represents the interquartile range (IQR). Adjusted p value (q) were computed by Benjamini-Hochberg method.

**Supplementary Fig.4** Single cell analysis of PBMC samples in COVID-19 patients and healthy donors. **a** tSNE plot displaying all identified cell types and states. **b** The markers distribution in cell types. The expression levels are blue color coded.

**Supplementary Fig.5** Single-cell based analysis in PBMC samples by sex. **a** Heatmap showed the z-score in male and female patients with critical COVID-19. **b** The expression of male-biased immune genes of squamous in the patients with critical COVID-19. Each dot means one cell, and the plot only show the genes positive expressed cells. For inside boxplots, the box represents the interquartile range (IQR). Adjusted p value (q) were computed by Benjamini-Hochberg method.

## Supplementary Tables

**Supplementary Table 1.** Statistics analysis of four COVID-19 outcomes across different age groups. (.xlsx).

**Supplementary Table 2.** The patient’s information and data sources of two single-cell RNA-sequencing datasets. (.xlsx).

**Supplementary Table 3.** Summary of gene-set enrichment analysis results of a nasal tissue-based single-cell RNA-sequencing dataset. (.xlsx)

**Supplementary Table 4.** Summary of gene-set enrichment analysis results of a PBMC-based single-cell RNA-sequencing dataset. (.xlsx)

**Supplementary Fig.1.**
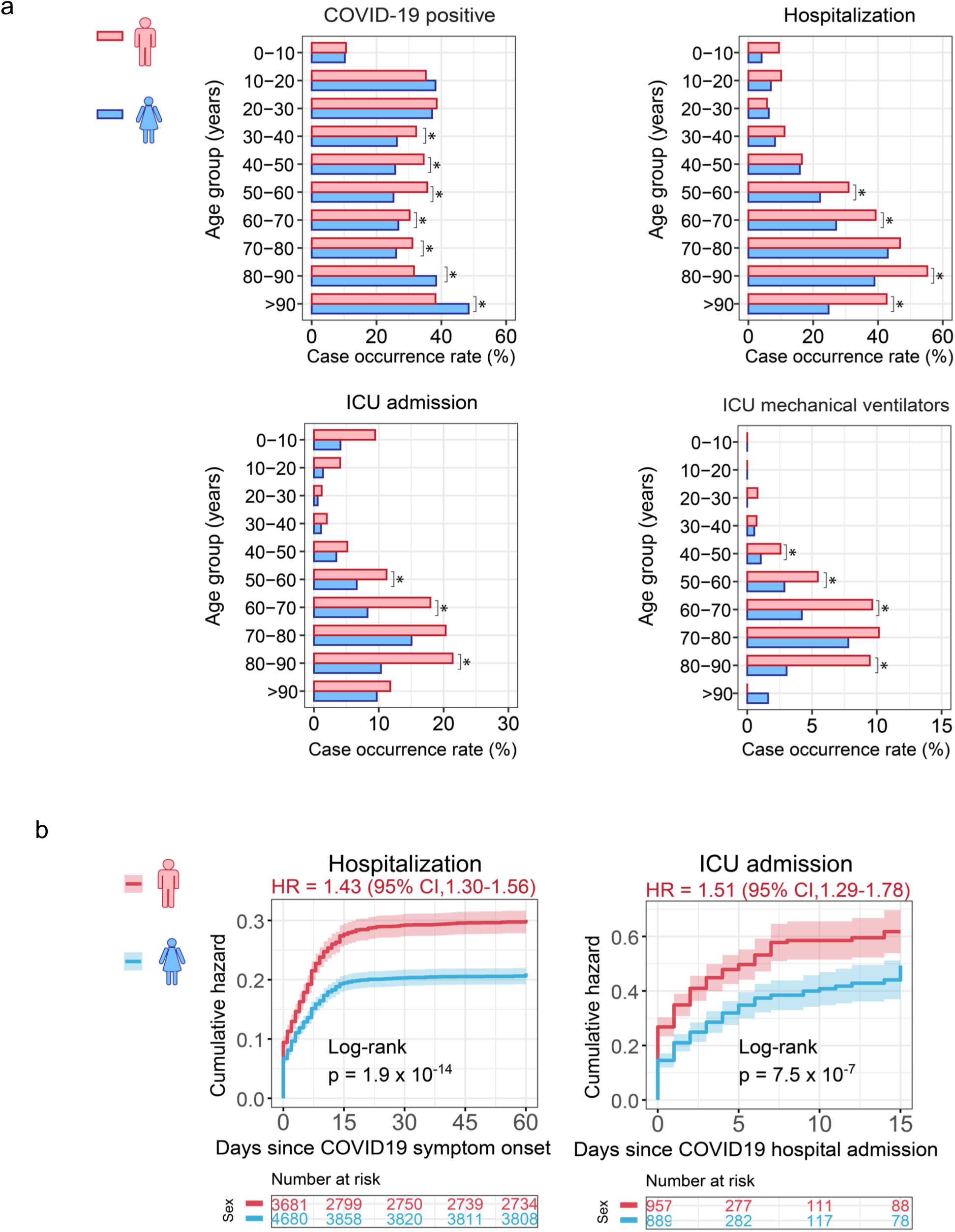
Clinical outcome and characteristics between male and female individuals with COVID-19 **a** Statistics analysis of four COVID-19 outcomes across different age groups. * denote p < 0.05 using Fisher extract test. **b** Cumulative hazard of hospitalization and ICU admission are shown. The results were computed using original cohort. The log-rank test with the Benjamini & Hochberg (BH) adjustment was used for comparing the statistical significance of cumulative hazard of hospitalization and ICU admission between men and women. The shadow represents 95% confidence interval. HR, hazard ratio.

**Supplementary Fig.2.**
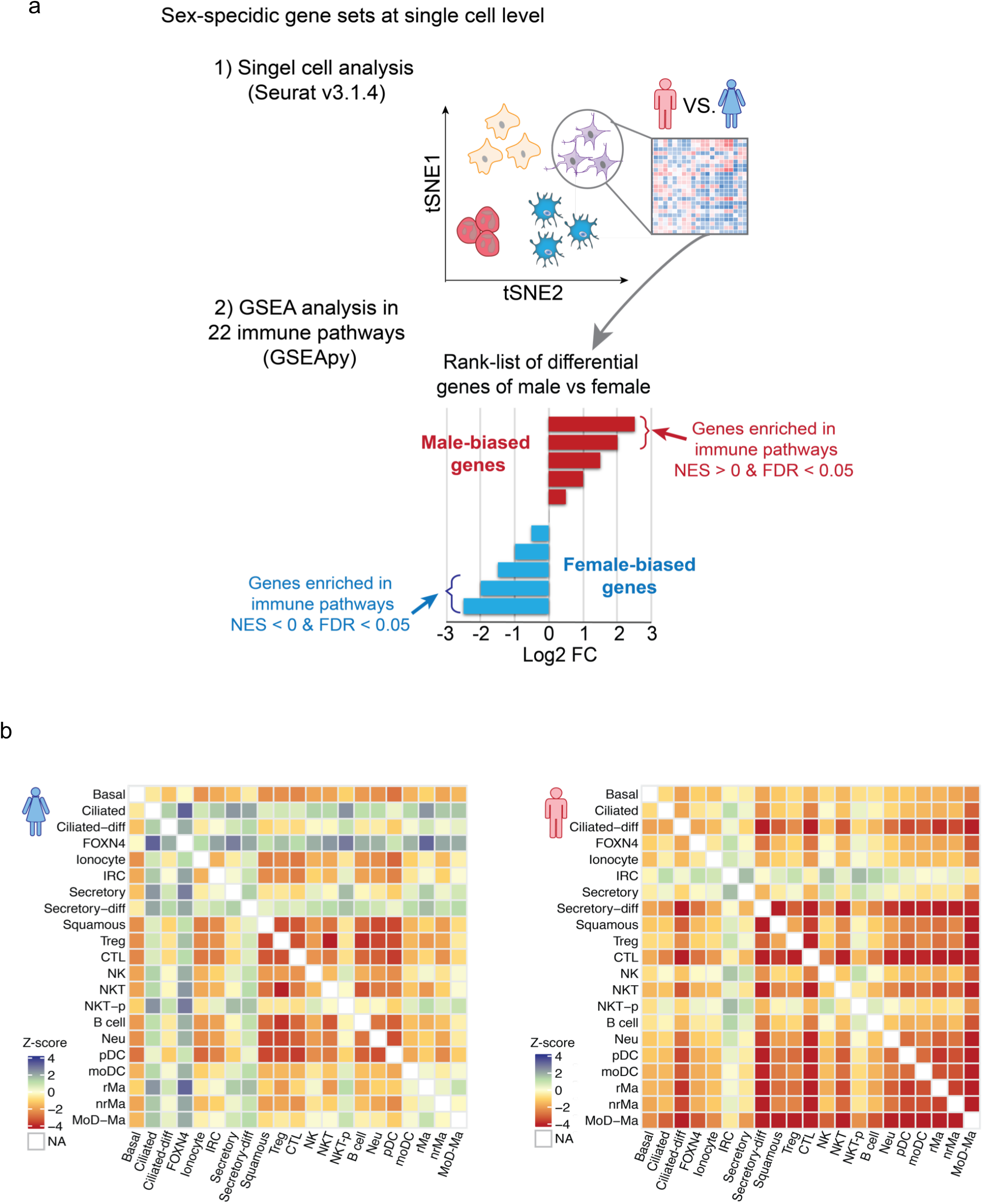
Cell types analysis of nasal samples by sex. **a** Workflow of GSEA analysis. **b** Heatmap showed the z-score in male and female patients with critical COVID-19.

**Supplementary Fig.3.**
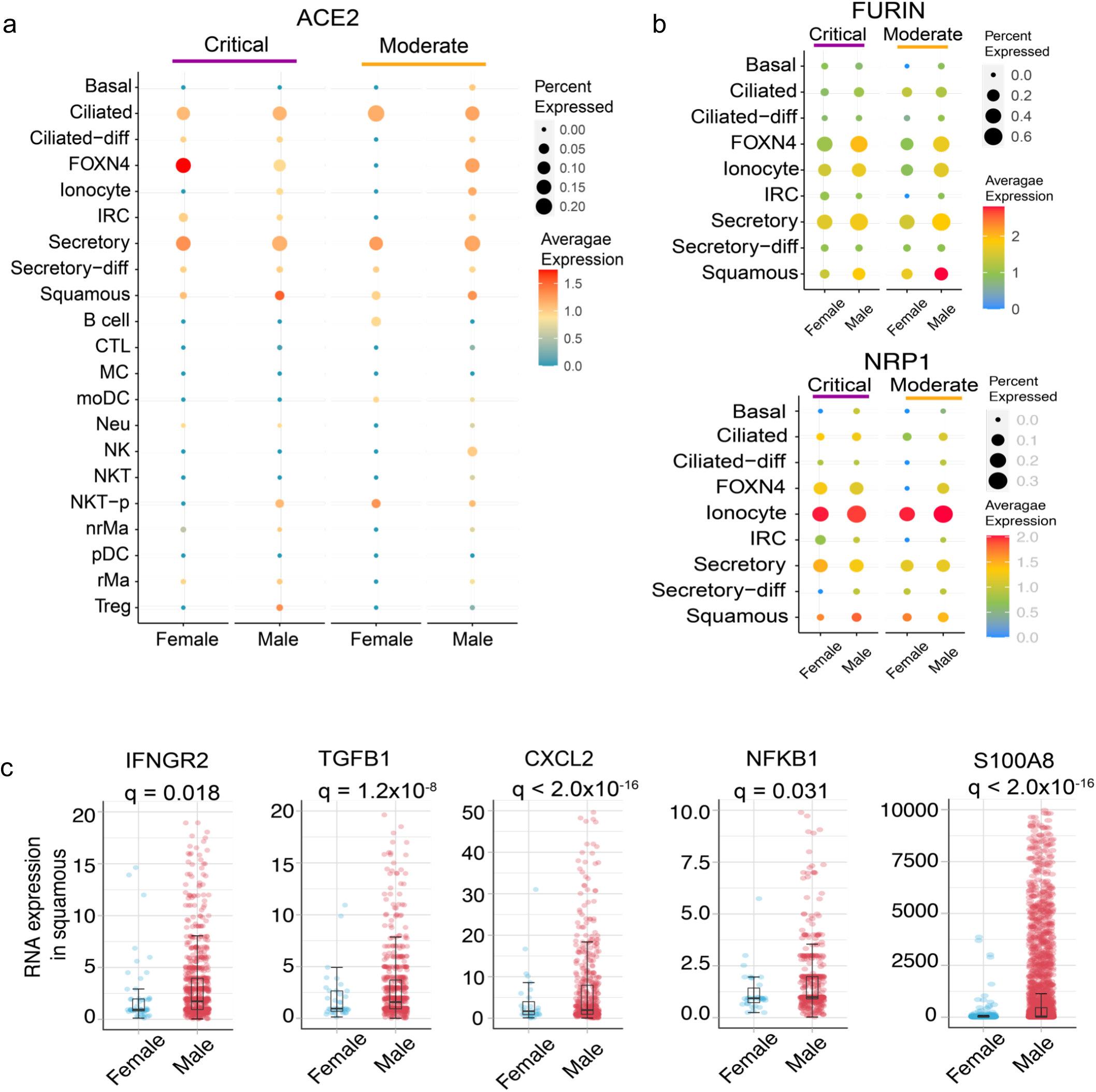
The expression of ACE2 and immune genes by sex. **a** ACE2 expression by sex in 22 cell types across critical and moderate COVID-19. conditions. The size of dot denotes the percentage of ACE2 or TMPRSSE positive expressed cells. The gradient color bar represents the average expression of genes in each cell type. **b** the dot plot showed the expression level and distribution of FURIN and NPR1 in epithelial cells by sex. **c** The expression of male-biased immune genes of squamous in the patients with critical COVID-19. Each dot means one cell, and the plot only show the genes positive expressed cells. For inside boxplots, the box represents the interquartile range (IQR). Adjusted p value (q) were computed by Benjamini-Hochberg method.

**Supplementary Fig.4.**
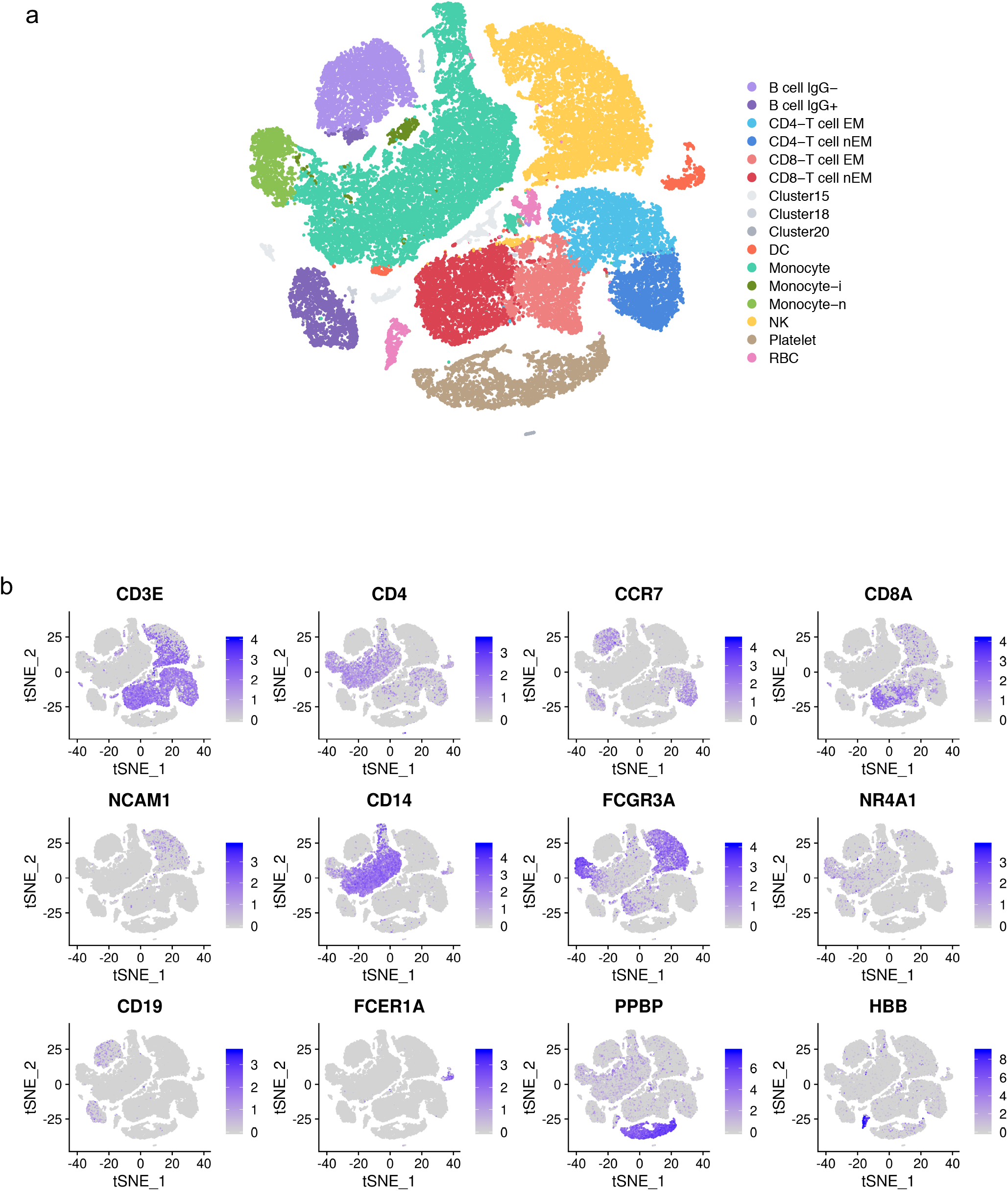
Single cell analysis of PBMC samples in COVID-19 patients and healthy donors. **a** tSNE plot displaying all identified cell types and states. **b** The markers distribution in cell types. The expression levels are blue color coded.

**Supplementary Fig.5.**
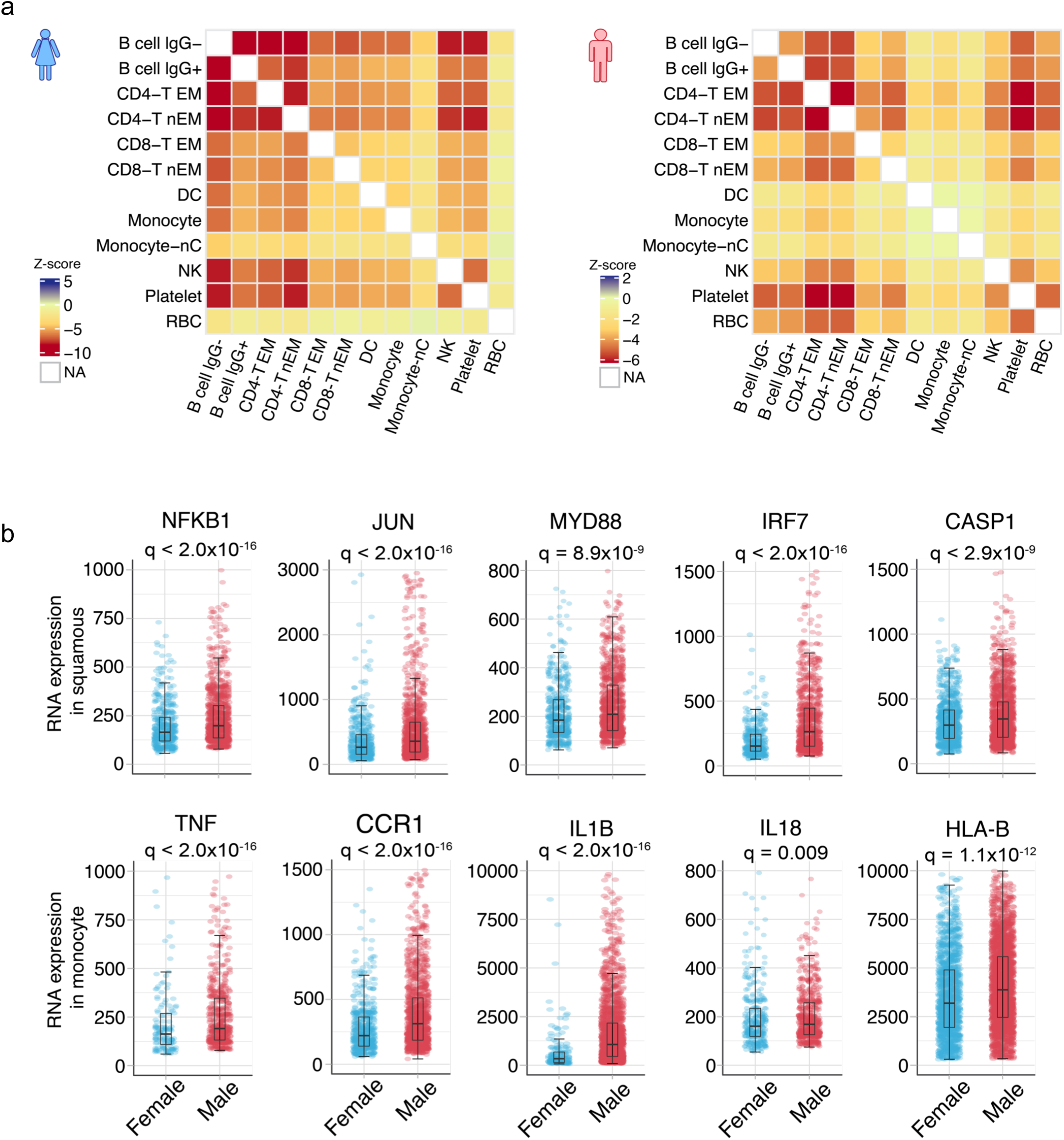
Single cell based analysis in PBMC samples by sex. **a** Heatmap showed the z-score in male and female patients with critical COVID-19. **b** The expression of male-biased immune genes of squamous in the patients with critical COVID-19. Each dot means one cell, and the plot only show the genes positive expressed cells. For inside boxplots, the box represents the interquartile range (IQR). Adjusted p value (q) were computed by Benjamini-Hochberg method.

